# Adult-born neurons maintain hippocampal cholinergic inputs and support working memory during aging

**DOI:** 10.1101/311423

**Authors:** Greer S. Kirshenbaum, Victoria K. Robson, Rebecca M. Shansky, Lisa M. Savage, E. David Leonardo, Alex Dranovsky

## Abstract

Adult neurogenesis is impaired in disorders of stress, memory, and cognition though its normal function remains unclear. Moreover, a systems level understanding of how a small number of young hippocampal neurons could dramatically influence brain function is lacking. We examined whether adult neurogenesis sustains hippocampal connections across the life span. Long-term suppression of neurogenesis as occurs during stress and aging resulted in a progressing decline in hippocampal acetylcholine and the slow emergence of profound working memory deficits. These deficits were accompanied by compensatory rewiring of cholinergic dentate gyrus inputs such that ventrally projecting neurons were recruited by the dorsal projection. Our study demonstrates that hippocampal neurogenesis supports memory by maintaining the septohippocampal circuit across the lifespan. It also provides a systems level explanation for the progressive nature of memory deterioration during normal and pathological aging and indicates that the brain connectome is malleable by experience.

## Introduction

Connectivity between brain systems is thought to be established during developmental critical periods and then remain relatively fixed in adults. However, neurogenesis persists in the hippocampus throughout life, raising a possibility that the addition of new cells could facilitate systems-level circuit rewiring over time. This is especially intriguing since adult neurogenesis is highly susceptible to the effects of environmental changes and aging (Knoth et al., 2010; Kuhn et al., 1996; Spalding et al., 2013) presenting a putative mechanism for a very slow form of experience-based circuit plasticity in the adult brain. Stress and aging-induced reductions in neurogenesis correlate with decreased cognitive (van Praag et al., 2005) and emotional flexibility (Snyder et al., 2011). However, little is known about how ongoing neurogenesis influences hippocampal circuitry and function over time.

The role of young hippocampal neurons has been closely examined. Several studies have elucidated the dynamic properties of new hippocampal neurons throughout their maturation, and others have illuminated how new neurons integrate into existing brain circuits (Zhao et al., 2008). Young, 4-12 week old, hippocampal neurons have been found to play a role in the more difficult versions of hippocampal dependent cognitive tasks (Clelland et al., 2009; Shors, 2008) and a subtle, but significant role in stress regulation (Snyder et al., 2011). These findings emerged from short-term experimental reductions or increases in neurogenesis in rodents and together indicate a modest, but significant contribution of young neurons to hippocampal function. However, neurogenesis is an ongoing process, raising the possibility that long-term changes in the addition of new neurons to existing circuits could result in system-wide changes with more dramatic consequences for hippocampal function and for behavior. This notion is especially intriguing since it entertains time scales that reflect aging and the chronicity of illnesses where neurogenesis and hippocampal function are impaired. We therefore hypothesized that long periods of reduced neurogenesis, as observed in aging and chronic stress, would influence hippocampal connectivity with other brain systems. Since new neurons are added throughout life, we expected that long periods of reduced neurogenesis could lead to substantial connectivity changes.

We investigated whether long term reductions in hippocampal neurogenesis influences inputs into the hilus where hippocampal neurogenesis occurs. We reduced neurogenesis for 5 months in adult mice and observed a slowly progressing reduction of hippocampal acetylcholine and impairments in working memory. These changes corresponded to a significant remodeling of the cholinergic septohippocampal projection with recruitment of ventrally projecting neurons for innervation of both the dorsal and ventral hilus.

## Results

### Long-term suppression of neurogenesis induces rewiring of cholinergic hilar inputs

We hypothesized that over time, reduction in neurogenesis would influence connectivity to the hilus, a structure where hippocampal afferents from many brain regions terminate to regulate neurogenesis, granule cell activity, and hippocampal function (Amaral et al., 2007). To model a reduction in hippocampal neurogenesis across aging, 2 month-old mice were treated with focal hippocampal X-irradiation (Santarelli et al., 2003) thereby dramatically and permanently reducing hippocampal neurogenesis (Figure S1). This method is thought to acutely target proliferating cells and also interfere with neural differentiation in the subgranular zone without dramatically influencing hippocampal and extra-hippocampal structures (Santarelli et al., 2003). We then used recombinant canine adenovirus (CAV), which undergoes selective retrograde neuronal transport (Kissa et al., 2002), to assess changes in hippocampal inputs over time. We injected CAV encoding green fluorescent protein (CAV-GFP) into the dorsal hilus of NG+ (Sham-irradiated mice with normal neurogenesis) and NG- (X-irradiated mice with diminished neurogenesis) animals after 5 or 2 months of reduced neurogenesis (Figure 1A). Following infusions of CAV-GFP, we observed cell bodies in numerous brain regions including the medial septum-nucleus of the diagonal band (MS-NDB) where the number of labeled cells appeared to be different in NG- mice after 5 months of reduced neurogenesis. MS-NDB was also particularly interesting since its hippocampal cholinergic projections are thought to regulate neurogenesis (Berg et al., 2013; Mohapel et al., 2005), be important for short-term memory (Chang and Gold, 2004), and exhibit deleterious changes during aging (Banuelos et al., 2013; Stroessner-Johnson et al., 1992). Compared to NG+ mice, NG- mice with a 5-month long reduction of neurogenesis had more GFP+ cells in the MS-NDB both ipsilateral and contralateral to the injection site (Figure 1B,C), indicating that additional MS-NDB neurons were projecting to the hilus in NG- animals. The change in the septohippocampal cholinergic inputs was not detectable in NG- mice with a 2 month-long reduction in neurogenesis (Figure 1C), indicating that this type of remodeling requires extended periods of living without neurogenesis. The change in connectivity was not detected in another major hilus projection, the locus coeruleus (LC; Figure 1B,C), indicating that reduction of neurogenesis causes changes in some, but not other hilar inputs. We wanted to verify that the circuit rewiring observed after 5 months with reduced neurogenesis was due to suppression of neurogenesis rather than an off-target effect of X-irradiation. We therefore used a genetic technique to suppress neurogenesis for 5 months in 2 month old mice and repeated our tracing experiments. New neurons generated in the subgranular zone develop from radial astrocytes that express glial fibrillary acidic protein (GFAP;(Garcia et al., 2004). We used mice that express herpes simplex virus thymidine kinase (Tk) under control of a GFAP promoter to suppress neurogenesis by administration of the antiviral drug valgancyclovir (VGCV) for 5 months. This approach was validated for targeting dividing stem cells and diminishing neurogenesis while sparing non-stem astrocytes (Garcia et al., 2004). After 5 months of VGCV treatment we observed suppression of neurogenesis in NG-^TK^ mice (GFAP-Tk+/− mice treated with VGCV for 5 months) compared to NG+^TK^ (GFAP-Tk-/- mice treated with VGCV for 5 months; Figure S1). We also observed that more cells bodies were labeled in the MS-NDB but not in the LC of NG-^TK^ mice compared to NG+^TK^ mice (Fig 1C) echoing the results observed in X-irradiated mice. Together the results demonstrate rewiring of septohippocampal inputs in animals with diminished neurogenesis.

**Figure 1.**
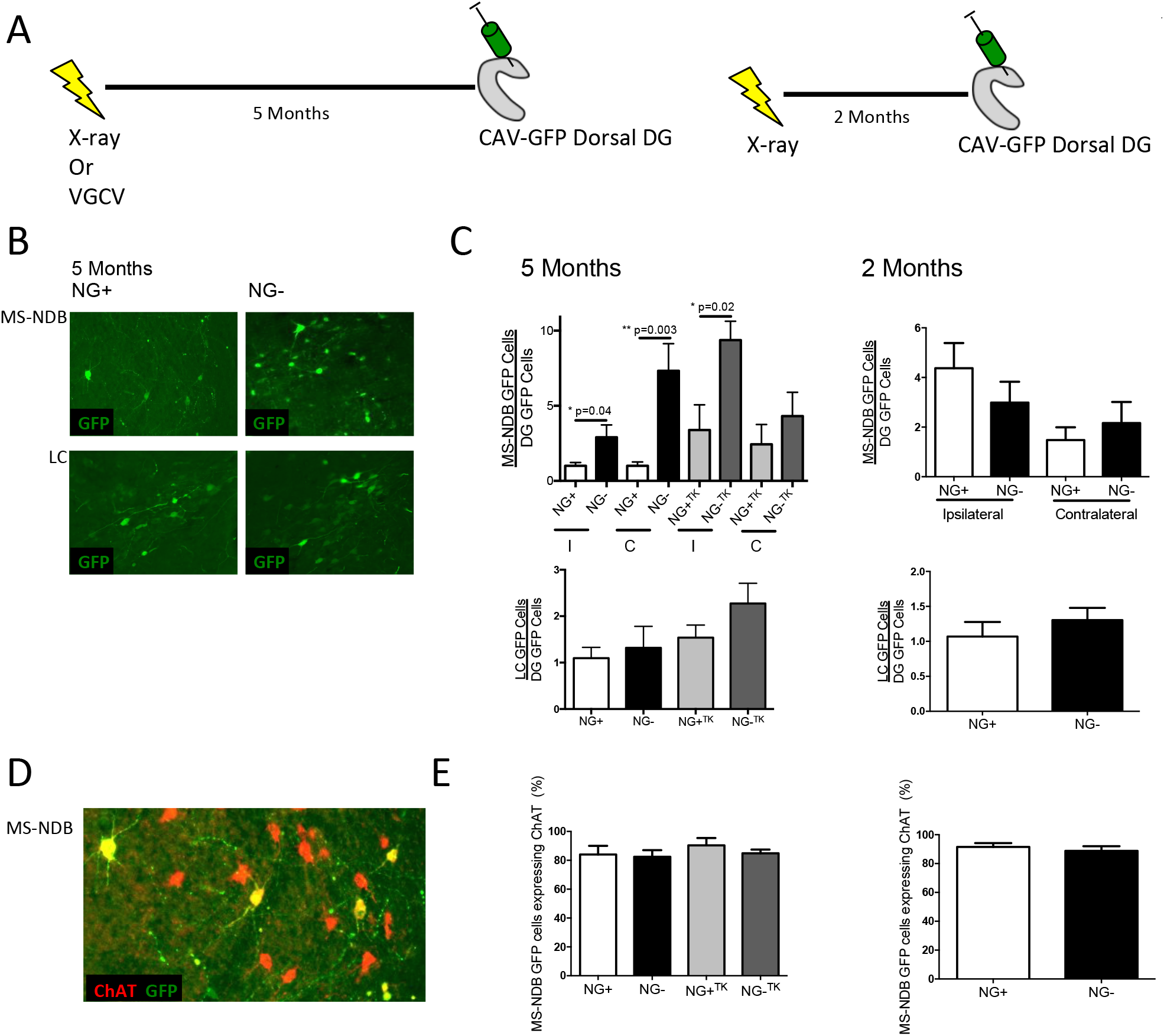
Increased cholinergic hilar inputs in mice living with diminished neurogenesis. (A) 2 month-old mice were exposed to focal hippocampal X-irradiation or VGCV. CAV-GFP was injected into the dorsal hilus 5 months after X-irradiation or VGCV treatment. CAV-GFP was injected also into a group of X-irradiated mice after two months. (B) GFP+ cells in the MS-NDB and the LC in NG+ and NG- mice (n=12 per group). (C) NG- mice without neurogenesis for 5 months showed significantly more cells projecting to the dorsal hilus than NG+ mice from the MS-NDB both ipsilateral (I) (t(22)=2.238, p= 0.0357) and contralateral (C; t(22)=3.316, p= 0.0033) to the injection site. NG- mice without neurogenesis for 2 months (n=5) and NG+ mice (n=5) showed similar connectivity from MS-NDB and LC to the dorsal hilus. NG-^TK^ mice with suppressed neurogenesis for 5 months showed significantly more cells projecting to the dorsal hilus than NG+^TK^ mice from the MS-NDB ipsilateral to the injection site (t(7)=2.927, p= 0.0221). (D, E) In the MS-NDB 80-90% of GFP labeled cells overlap with a primary marker of cholinergic cells, acetylcholineserase in 5 and 2 month groups. I= Ipsilateral, C= Contralateral. Bars represent mean ± SEM.

The MS-NDB hippocampal projection is comprised of roughly two thirds GABAergic, one third cholinergic and a small proportion of glutamatergic neurons (Amaral et al., 2007; Colom et al., 2005; Henderson et al., 2010). All three cell types project to the hilus (Amaral et al., 2007; Colom et al., 2005; Henderson et al., 2010). Remarkably, 80-90% of CAV-GFP labeled MS-NDB cells in NG+, NG-, NG+^TK^and NG-^TK^mice co-expressed the cholinergic marker choline acetyltransferase (ChAT), but not markers for the dominant GABAergic populations residing in the region (Figure 1 D-E, Figure S2). Our results therefore signify that CAV primarily transduces cholinergic hilar inputs. In the autonomic nervous system mature neurons are occasionally thought to convert to a ChAT expressing phenotype (Apostolova et al., 2010). Moreover, recent reports suggest that CNS neurons can begin to express new identity markers in adult animals (Dulcis et al., 2013). We therefore also examined if the total number of ChAT+ cells was changed in animals living without neurogenesis. We found that the total number of cholinergic neurons in the MS-NDB was not different in NG+ and NG- mice or NG+^TK^and NG-^TK^ mice (Figure S3), suggesting that the increase in labeled cells was due to recruitment of inputs into the region from already existing cholinergic neurons that project elsewhere in NG+ mice.

### Ongoing neurogenesis maintains working memory

Aging and chronic stress reduce neurogenesis (Zhao et al., 2008) while compromising cholinergic function (Nyakas et al., 2011; Terry and Buccafusco, 2003), and short-term memory (Zhao et al., 2008). Therefore, an increase in cholinergic inputs as a result of neurogenesis ablation seemed surprising. We therefore hypothesized that increased inputs in our models of reducing neurogenesis across aging reflected a natural compensation for an evolving impairment in cholinergic function. We tested NG- mice in a 4-arm spontaneous alternation task, which is dependent on an intact cholinergic septohippocampal projection(Chang and Gold, 2004). We found that NG+ and NG- mice performed at similar levels when neurogenesis was reduced for 2 and 4 months. However, a deficit in short-term memory emerged in NG- mice after 5 months, coinciding with changes in the septohippocampal projection (Chang and Gold, 2004). This deficit remained after 12 months without neurogenesis, when NG- animals performed at chance levels demonstrating a failure of working memory in this task (Figure 2 A-E). Thus, a reduction of neurogenesis has a slowly emerging, but profound effect on a cholinergic-dependent working memory task.

**Figure 2.**
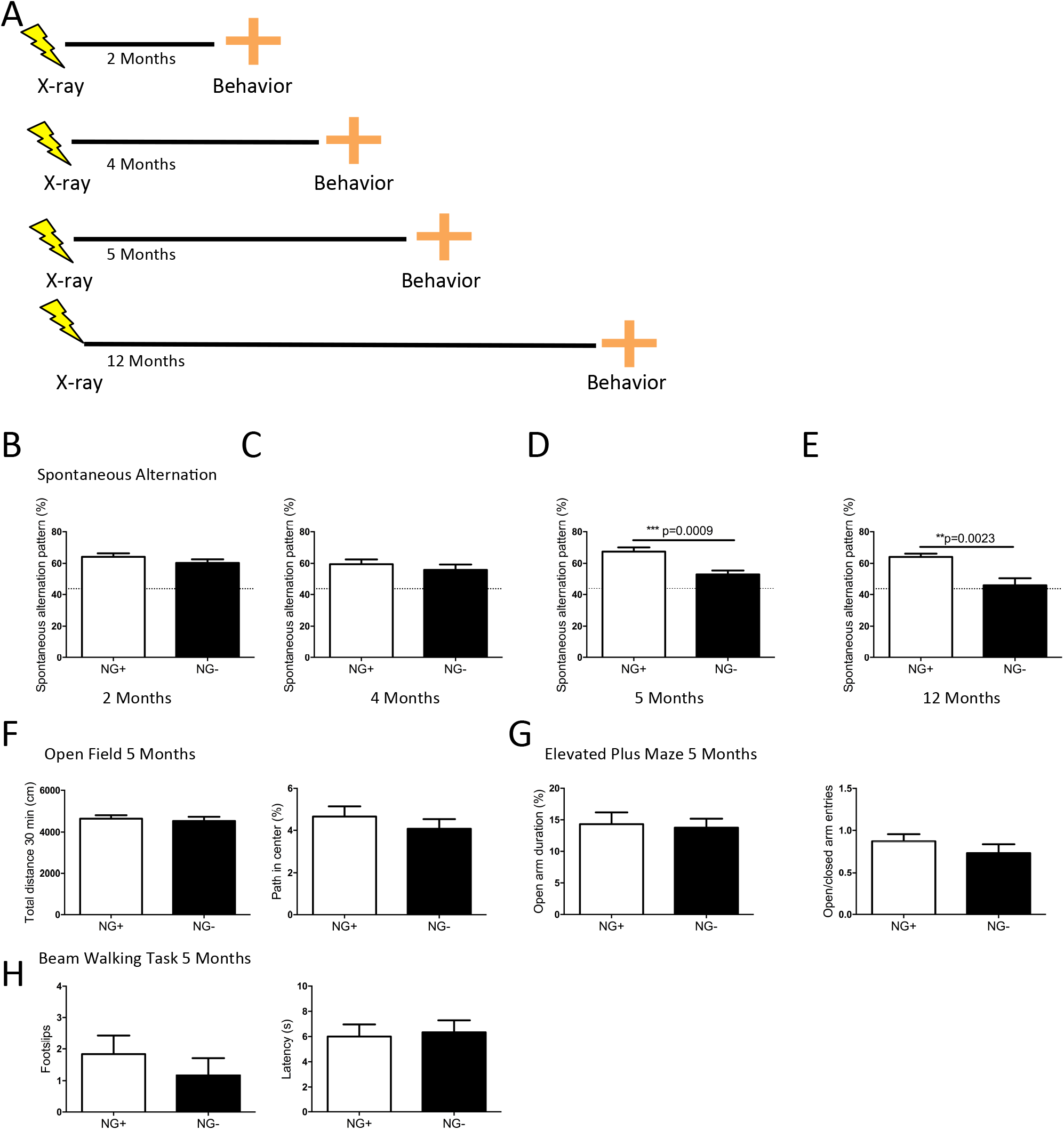
A working memory deficit emerges in mice after a prolonged reduction of adult neurogenesis. (A) Behavior of NG- mice was assessed at 2, 4, 5 and 12 months without neurogenesis and compared to age matched NG+ controls. (B-E) Spontaneous alternation pattern (SAP) in a 4-arm spontaneous alternation task. (B,C) NG- mice after 2 and 4 months without neurogenesis had similar SAP scores of about 60% as NG+ mice (2 months n=15 per group, 4 months NG+ n=12 NG- n=13). (D) However, after 5 months without neurogenesis a deficit emerged in NG- mice where they showed SAP scores around 50% (NG+ n=10, NG- n=14; t(22)=3.835, p= 0.0009). (E) After 12 months without neurogenesis SAP scores declined to chance levels of about 44% (dotted line) NG- mice (NG+ n=7, NG- n=5; t(10)=4.06, p= 0.0023). (F) In the open field, NG- mice (n=11) without neurogenesis for 5 months and NG+ mice (n=8) showed no differences in total distance travelled in 30 min (cm) or the percentage of time spent in the center. (G) In the elevated plus maze NG- mice (n=8)without neurogenesis for 5 months and NG+ mice (n=11) showed similar open arm duration (s) and similar ratios of open to closed arm entries. (H) In a beam walking task NG- mice (n=6) without neurogenesis for 5 months and NG+ mice (n=6) showed a similar number of total footslips and traversal latency (s). Bars represent mean ± SEM.

Several learning and anxiety tasks have been examined in mice with short-term ablation of hippocampal neurogenesis(Drew et al., 2010). We confirmed reports that NG- mice have an impairment in a form of contextual fear conditioning after 8 weeks without neurogenesis, but display unimpaired anxiety-related behavior in the open field or elevated plus maze (Figure S4 and Figure 2F,G) neither of which depend on cholinergic functioning or neurogenesis. Importantly, NG- animals were not impaired on a beam-walking task (Figure 2H), which requires an intact cholinergic nucleus basalis projection to the frontoparietal cortex (Lehmann et al., 2002). This demonstrated that the reduction in neurogenesis alters the septohippocampal projection system, but not other cholinergic circuitry.

### Cholinergic dysfunction precedes input rewiring and memory deficits

Given the delayed emergence of our findings, we hypothesized that changes in septohippocampal circuit function precede our behavioral and anatomic observations. We tested this by challenging the cholinergic system in NG- mice after 2 months without neurogenesis with the muscarinic cholinergic antagonist scopolamine, which impairs spontaneous alternation behavior in high doses (Andriambeloson et al., 2014). As expected, at a normally sub-threshold dose, scopolamine did not affect NG+ mice, however it profoundly impaired spontaneous alternation behavior in NG- mice even after 2 months without neurogenesis (Figure 3A). This experiment indicates that cholinergic dysfunction is present in NG- animals long before behavioral changes emerge. It also supports the hypothesis that changes in septohippocampal inputs observed in Figure 1 may be compensatory to a primary loss in cholinergic tone. Since we observed no decrease in septal cholinergic neurons projecting to the hippocampus (Figure 1), we hypothesized that deficits in spontaneous alternation could be rescued by bolstering the endogenous cholinergic tone in NG- mice. We administered physostigmine, a cholinesterase inhibitor, which increases synaptic acetylcholine, to NG- mice and found that the spontaneous alternation deficit could be acutely and fully restored even after 12 months without neurogenesis (Figure 3B). Together these results strongly suggested a slowly emerging acetylcholine deficit in NG- mice.

**Figure 3.**
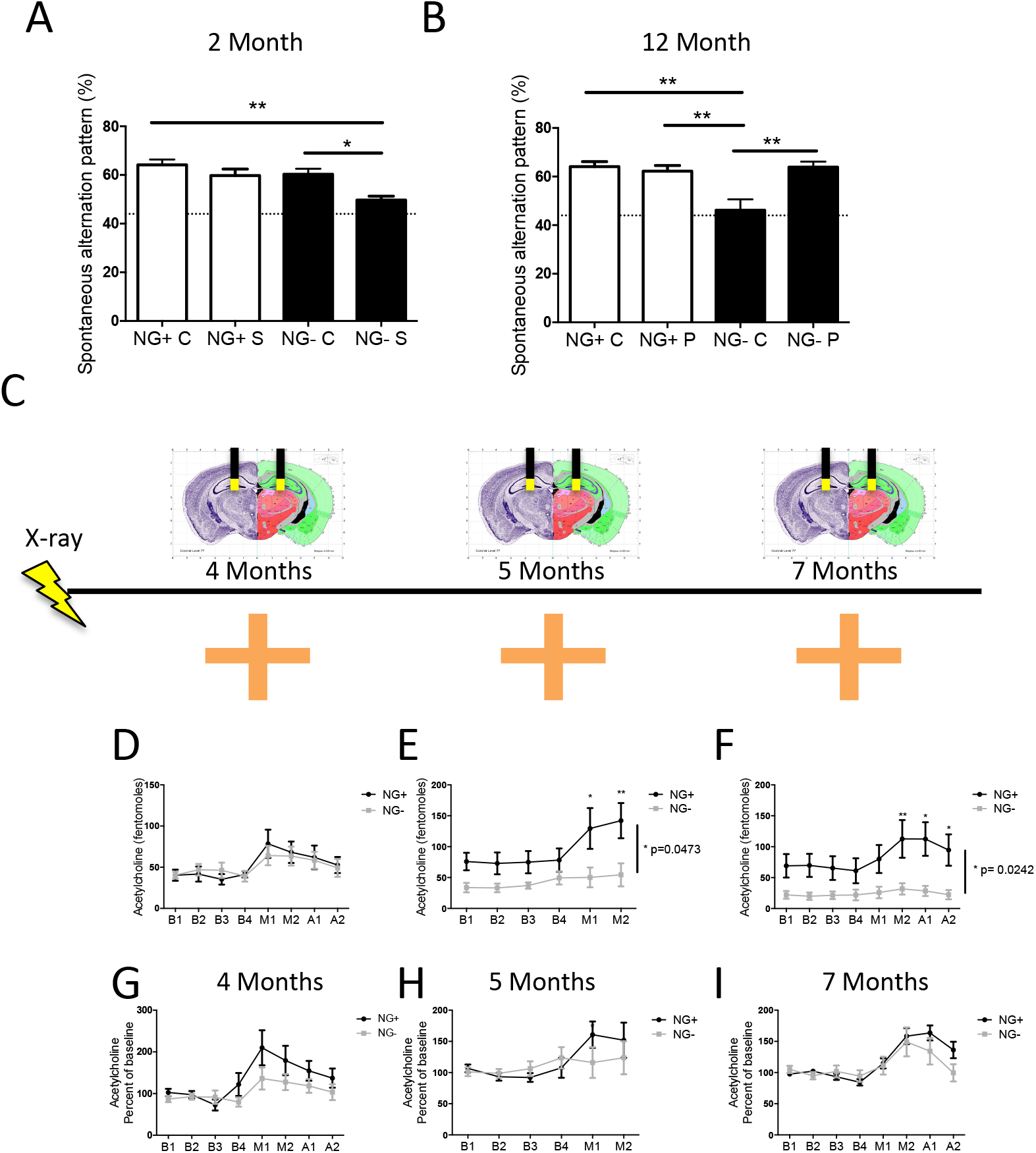
Hippocampal acetylcholine release declines in mice after a prolonged reduction of adult neurogenesis. (A) NG- mice without neurogenesis for 2 months and NG+ mice were administered a muscarinic acetylcholine receptor antagonist scopolamine and spontaneous alternation pattern was assessed in a 4-arm spontaneous alternation task. Performance was compared across all groups (NG+ C n= 15, NG- C n=15, NG+ S n=5 and NG- S n=6, main effect of group F_337_ = 4.977, p= 0.0053). The performance of NG- S mice was reduced compared to all groups to near chance levels (dotted line). (B) NG- mice without neurogenesis for 12 months and NG+ mice were administered an acetlycholinesterase inhibitor physostigmine and spontaneous alternation pattern was assessed in a 4-arm spontaneous alternation task. Performance was compared across all groups (NG+ C n= 7, NG- C n=5, NG+ P n=7, NG- P n=5, main effect of group F_3,26_ = 8.683, p= 0.0007). NG- C mice performed below all groups and at chance. Bars represent mean ± SEM, Tukey’s *post hoc* * p<0.05, ** p<0.01. (C) Mice were treated with X-irradiation or Sham treatment and bilaterally implanted with cannulae. In one group of mice microdialysis measurements were taken from the left dorsal hilus after 4 months without neurogenesis. In a separate group of mice microdialysis measurements were taken from the right dorsal hilus at 5 months and the left dorsal hilus at 7 months without neurogenesis. Microdialysis measurements were taken at baseline (B1-B4), while mice were in the 4-arm spontaneous alternation maze (M1-M2) and after being removed from the maze (A1-A2). (D) After 4 months without neurogenesis NG+ and NG- mice showed similar patterns of acetylcholine release that increased when animals were in the maze (NG+ n= 7, NG- n=8; main effect of time F_791_ = 8.504, p< 0.0001). (E) After 5 months without neurogenesis NG- mice showed significantly lower acetylcholine release (n=6 per group; main effect of group F_110_ = 5.109, p= 0.0473), main effect of time F_550_ = 10.87, p< 0.0001, interaction effect F_550_= 4.198, p= 0.0029, (F) Reduced acetylcholine in NG- mice remained after 7 months with reduced neurogenesis (NG+ n= 5, NG- n=6; main effect of group F_19_ = 7.31, p= 0.0242, main effect of time F_763_ = 13.44, p< 0.0001, interaction effect F_763_= 6.904, p< 0.0001). (G) Mice without neurogenesis for 4 months demonstrated an increase in acetylcholine above baseline during the spontaneous alternation task. NG- mice showed a slightly attenuated increase in acetylcholine while performing spontaneous alternation compared to NG+ controls (main effect of time F_791_= 8.921, p< 0.0001, main effect of group F_791_ = 1.874, p=0.1942, time × group interaction F_791_ = 1.786, p= 0.0996) (H) Mice without neurogenesis for 5 months demonstrated an increase in acetylcholine above baseline during the spontaneous alternation task (main effect of time F_540_ = 3.695, p = 0.0076) as did animals after 7 months without neurogenesis (main effect of time F_763_ = 10,68, p < 0.0001) (I). Bars and points represent mean ± SEM, Bonferroni *post hoc* *p<0.05, ** p<0.01.

We therefore used awake-behaving microdialysis and HPLC to directly measure hippocampal acetylcholine efflux both at baseline and during spontaneous alternation in mice with neurogenesis reduced for 4, 5 and 7 months (Figure 3C). We observed that only a prolonged reduction in neurogenesis lead to reduced acetylcholine in the DG. Four months without neurogenesis resulted in no change in DG acetylcholine levels at baseline or during spontaneous alternation (Figure 3D). However, after 5 and 7 months without neurogenesis, NG- mice exhibited reduced DG acetylcholine both at baseline and during spontaneous alternation (Figure 3E,F). Moreover, both NG+ and NG- groups at 4, 5 and 7 months mounted an increase in acetylcholine efflux while performing the spontaneous alternation task (Figure 3G-I), indicating that the cholinergic projection was engaged but the total release was attenuated. Interestingly, the 4 month group had a slightly attenuated acetylcholine increase while performing spontaneous alternation (Figure 3G). Together the results indicate that reduction in neurogenesis initially results in a subthreshold acetylcholine deficit, which is followed by attenuated release of phasic acetylcholine, progressing to a tonic efflux deficit and ultimately functional circuit deficits with compensatory changes in cholinergic innervation. Our findings therefore show that ongoing neurogenesis functions to maintain septohippocampal cholinergic tone and function.

### Ventral cholinergic projection is recruited by the dorsal hilus during rewiring

Normally, the septohippocampal projection has a highly organized topography. In rats, MS-NDB neurons project to either the dorsal or ventral hilus, but not to both (Ohara et al., 2013), though this has not been previously established in mice. We hypothesized that the increase in dorsally projecting MS-NDB cells in NG- mice after 5 months results from recruitment of cholinergic fibers of passage that normally traverse the dorsal to innervate the ventral DG. We analyzed the topography of MS-NDB cells projecting to the dorsal hilus in NG+ and NG- mice. We found that the increase in labeled cells in NG- mice after 5 months with reduced neurogenesis originated from structures that normally project to the ventral hippocampus, the medial NDB and lateral aspect of the MS in rats (Figure 4B). This suggested that, aging with reduced neurogenesis results in rewiring within the septohippocampal projection. We then directly tested our hypothesis by injecting a CAV-GFP into the dorsal hilus and a red reporter CAV (CAV-Cherry) into the ventral hilus in NG+ and NG- mice after 5 months of reduced neurogenesis (Figure 4C). We found that in NG+ mice, as in rats, dorsally projecting (GFP-labeled) cells were mostly distinct from ventrally projecting (Cherry labeled) cells (Figure 4D). However, in NG- mice «40% of cells with terminals in the dorsal hilus also had terminals in the ventral hilus (GFP and Cherry labeled; Figure 4D). This result indicates a rewiring of the septohippocampal projection in NG- mice where ventrally projecting MS-NDB cholinergic neurons also develop dorsal projections. Moreover it appears that ventrally projecting MS-NDB cholinergic neurons also demonstrated rewiring within the ventral hippocampus. NG- mice showed an increased number of Cherry labeled cells in the MS-NDB, specifically in the ventrally projecting medial NDB and lateral MS compared to NG+ mice (Figure 4E). Therefore, aging without neurogenesis results in increased innervation, albeit dysfunctional, in both the ventral and the dorsal hippocampus by normally ventrally projecting MS-NDB neurons. The findings provide a mechanism for increased numbers of septal cholinergic inputs into the dorsal hippocampus of NG- mice.

**Figure 4.**
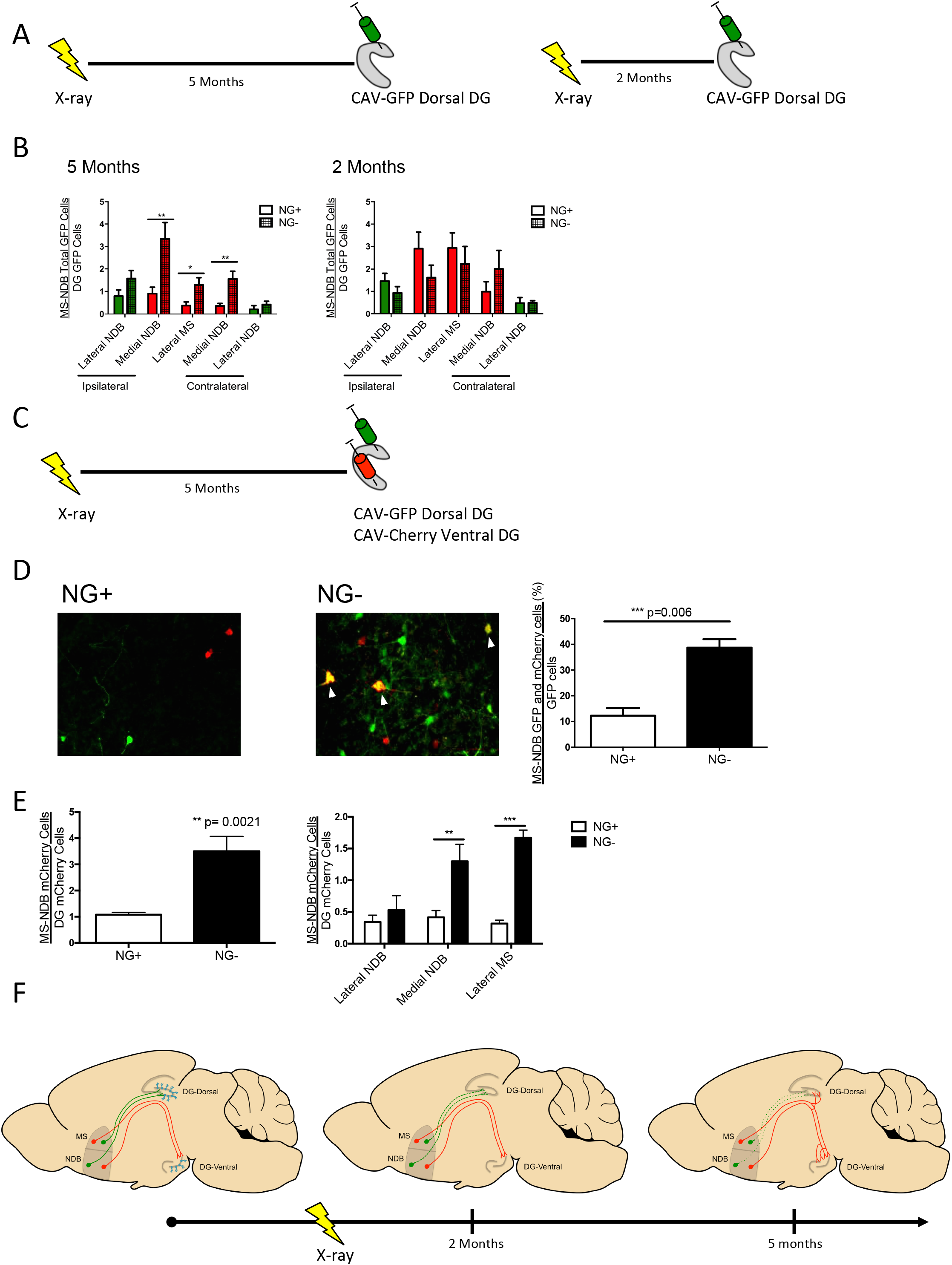
Rewiring of cholinergic septotemporal projection after a prolonged reduction in adult neurogenesis. (A) Mice were treated with X-irradiation or Sham treatment and CAV-GFP was injected into the dorsal hilus after 5 or 2 months. (B) After 5 months without neurogenesis, the dorsal hilus of NG- mice shows an increase in innervation by GFP+ cell bodies that are expected to primarily innervate the ventral hilus (Ohara et al., 2013) compared to NG+ mice (n=12 per group). NG+ mice show an increase in projections to the dorsal hilus from the medial NDB, contralateral (t(22)=3.171, p= 0.0044) and ipsilateral (t(22)=3.346, p= 0.0029) to the injection site, as well as the lateral MS (t(22)=2.554, p= 0.0181). This change has not emerged in NG- mice without neurogenesis for 2 months. (C) In NG+ and NG- mice without neurogenesis for 5 months, CAV-GFP was injected into the dorsal hilus and CAV-Cherry was injected into the ventral hilus. (E) In NG+ mice (n=5) 10% of GFP+ cells in the MS-NDB showed Cherry labeling and this significantly increased to 40% in NG- mice (n=4; t(7)=5.985, p= 0.0006). (E) NG+ mice (n=5) show a greater number of cells projecting from the MS-NDB to the ventral hilus compared to NG- mice (n=4, t(7)=4.754, p= 0.0021). The increase in cell bodies is in the medial NDB (t(7)=3.331, p= 0.0126) and lateral MS (t(7)=11.14, p<0.0001). Bars represent mean ± SEM. * p<0.05, ** p<0.01 (F) Structural and functional reorganization of the septohippocampal circuit in NG- mice. NG+ mice show acetylcholine release that supports working memory and cholinergic afferent organization within the septohippocampal projection. NG- mice without neurogenesis for 2 months show an emerging deficit in acetylcholine release in the hippocampus but maintain cholinergic afferent organization within the septohippocampal projection. NG- mice without neurogenesis for 5 months show significant reductions in hippocampal acetylcholine release and rewiring of cholinergic septohippocampal inputs with septal neurons that normally project to the ventral hilus innervating the dorsal hilus and increased innervation of the ventral hilus.

## Discussion

Our findings demonstrate a critical role for ongoing neurogenesis over extended periods of time in the maintenance of cholinergic hippocampal inputs and non-reinforced working memory. This represents a remarkably slow form of plasticity emerging over months. A temporal scale of such a magnitude corresponds to cumulative changes observed during chronic disease and aging. Accordingly, hippocampal neurogenesis is reliably reduced during chronic psychosocial stress, aging, and in Alzheimer’s disease (Perry et al., 2012; Shors, 2008). Here, we have observed that an experimental chronic reduction in neurogenesis is sufficient to result in systems level hippocampal dysfunction over time.

The earliest evidence of subthreshold cholinergic deficits in our study corresponds to time points when numerous, more subtle behavioral deficits associated with ablation of neurogenesis have been described (Clelland et al., 2009; Drew et al., 2010; Snyder et al., 2011). Our findings therefore raise a possibility that mild cholinergic dysfunction may underlie the more subtle behavioral deficits more proximately following ablation of young neurons. Studies using physostygmine could be used to address this possibility.

Ultimately, a profound degradation of DG-dependent working memory emerged in NG- mice along with a dramatic remodeling of septohippocampal cholinergic DG inputs. Septal cholinergic neurons were previously demonstrated to undergo axonal sprouting in the absence of synaptic targets (Stanfield and Cowan, 1982) and temporary remodeling while animals are housed in enriched environments (Bergami et al., 2015). Our findings indicate that elimination of adult-born neurons results in compensatory rewiring of the septohippocampal projection by recruitment of ventrally projecting axons for dual (dorsal-ventral) innervation. Thus, young dentate granule neurons appear to maintain the integrity of the septohippocampal circuit over the lifespan. We hypothesize that hippocampal neurogenesis serves as a functional target for cholinergic septohippocampal neurons, and in their absence cholinergic cells initially show a reduction in cholinergic activity, then reorganize their innervation as a compensatory mechanism (Figure 4F).

The progressive nature of behavioral and anatomic changes presented here is reminiscent of neurodegenerative changes observed during age-related cognitivedecline and in Alzheimer’s disease, where decreased cholinergic innervation was previously linked to fewer neural stem cells (Terry and Buccafusco, 2003). Hence, neurogenesis may constitute a target for stabilizing the septohippocampal circuit and consequently working memory during normal aging and in disease states. More broadly, given the susceptibility of adult neurogenesis to the effects of chronic stress and aging, it is intriguing to speculate that the adult connectome is more malleable by experiences than currently appreciated. Our results raise the possibility that targeting distinct populations of cells constitutes a viable strategy for rewiring of long-range connectivity in the brain.

## Author Contributions

Conceptualization, GSK and AD; Methodology GSK, LMS and AD; Validation, GSK. Formal analysis, GSK; Investigation, GSK, VKR, RMS and LMS; Resources, GSK; Data Curation, GSK; Writing – Original draft GSK and AD; Writing - Review and Editing, GSK, EDL and AD; Visualization, GSK, VKR and AD; Supervision EDL and AD; Project administration, GSK; Funding Acquisition, GSK, LMS, EDL and AD.

## Acknowledgements

The authors thank Joshua Gordon and Steven Siegelbaum for critical reading of the manuscript and members of the ADL and Gordon labs for helpful insights. GSK was supported by a Canadian Institutes of Health Research Postdoctoral Fellowship. This work was supported by MH091844 and NARSAD young investigator award (AD), MH091427 (EDL), and NS085502 (LMS). AD is an Irving Scholar. Authors declare no conflicts of interest.

## Experimental Procedures

### Animals

Focal X-ray treatment of the hippocampus leads to long-term reductions of cell proliferation in the dentate gyrus and spares irradiation to most of the brain(Santarelli et al., 2003). Male C57BL/6J mice (Jackson Laboratories) were used in experiments of NG+ (sham) and NG- (X-irradiated) mice. Mice were delivered to our animal facility at 7-weeks-old and acclimated for one week before sham or X-irradiation treatment at 8 weeks.

GFAP-Tk heterozygous mice (NG-^TK^; (Bush et al., 1998) express herpes thymidine kinase (Tk) under control of the glial fibrillary acidic protein (GFAP) promoter, expressed in stem and nonstem astrocytes. Treatment with Valgancyclovir (VGCV) in NG-^TK^ mice leads to reductions in cell proliferation in dividing stem cells with relative sparing of non-stem astrocytes (Garcia, 2004). Female GFAP-Tk heterozygous mice were mated with wild-type littermate males, all on C57BL/6J 129S6 mixed background. Male pups were genotyped using previously reported PCR reactions (Bush et al., 1998), weaned at P21, and housed 3–5 per cage with mixed genotypes. Half the mice were GFAP-Tk heterozygous (NG-^TK^) and half were negative for the gene (NG+^TK^). At 8 weeks mice were started on a feeding schedule of chow containing VGCV.

Mice were given *ad libitum* access to food and water under a 12:12 h light:dark cycle in a temperature-controlled (72°F) colony. All animal experiments were performed in accordance with the Guide for the Care and Use of Laboratory Animals and approved by the New York State Psychiatric Institute Animal Care and Use Committee.

### X-irradiation

Similar to previous studies(Santarelli et al., 2003), 8-week-old mice were anesthetized with ketamine and xylazine (150 mg/kg and 10 mg/kg respectively). NG+ mice were untreated. NG- mice were placed in a stereotaxic frame, covered by a lead shield with a 3.22 × 11 mm opening over the hippocampus (interaural 3.00 to 0.00) and placed in a X-RAD 320 biological irradiator (PXI; North Branford, CT). The X-RAD 302 operated at 300kV and 12 mA with a 2-mm AI filter and delivered 2.5-Gy doses per X-ray session. Mice were treated for three sessions, separated by a 2 day interval (day 1, 4 and 7) so NG- mice received a total dose of 7.5-Gy. To assess the effect of X-ray on cell proliferation, tissue from NG+ and NG- mice were immunolabeled for a marker for cell division Ki-67 and a marker of immature neurons DCX.

### Valgancyclovir treatment in GFAP-Tk^+/−^ Mice

Chow containing VGCV (165 mg/kg) was administered to mice at 8 week until 5 months of age. Mice were on a feeding schedule of chow where they were fed VGCV chow for 5 days and normal chow for 2 days. This feeding schedule was employed to reduce gastrointestinal side effects caused by Tk expression in gut tissues. To assess the effect of VGCV on cell proliferation, tissue from GFAP-Tk^-/-^ (NG+^TK^) and GFAP-Tk^+/−^ (NG-^TK^) mice were immunolabeled for a marker for cell division Ki-67 and a marker of immature neurons DCX.

### Retrograde Tracing

To visualize cell bodies projecting to the dorsal dentate gyrus, we used a canine adenovirus, expressing GFP (CAV-GFP), which is taken up by axon terminals and transported to cell bodies(Kissa et al., 2002). NG+ and NG- mice at 5 or 2 months after treatment were used.

Mice were anesthetized with a ketamine, xylazine, acepromazine mixture (65mg/kg, 13mg/kg, 1.5mg/kg respectively) and placed into a stereotaxic frame (David Koph Instruments) with the skull exposed. A 10μl Hamilton syringe with pulled glass pipette was used to infuse 1.5μl of CAV-GFP (5 × 10^12^) to the right dorsal hilus (bregma coordinates: anteroposterior -2.3mm, mediolateral 1.6mm, dorsoventeral -1.6mm) at 0.2 μl/min. Mice were sacrificed 1-4 weeks following surgery.

To visualize cell bodies projecting to the dorsal hippocampus and the ventral hippocampus, we infused CAV-GFP in the dorsal hippocampus and CAV-cherry in the ventral hippocampus. A group of 7-month-old NG+ and NG- mice were infused with CAV-GFP in the right dorsal hilus as described above. In addition the right ventral hilus (bregma coordinates: anteroposterior - 3.2mm, mediolateral 2.3mm, dorsoventeral -4.3mm) was infused with 1.5 μl of CAV-cherry (5 × 10^12^) at 0.2 μl/min. Mice were sacrificed 1-4 weeks following surgery.

### Immunohistochemistry

Mice were anesthetized with a ketamine and xylazine mixture (150mg/kg and 10mg/kg respectively) and transcardially perfused with ice cold phosphate-buffered saline (PBS; pH 7.4) followed by 4% paraformaldehyde (PFA) in PBS. Brains were stored in 4% PFA overnight and transferred to 30% sucrose for 48h. Brains were sagittally sectioned at 35 μm and stored in PBS with 0.02% azide.

For immunostaining tissue was washed in PBS, blocked with 10% normal donkey serum and incubated in primary antibody overnight at 4°C. The following primary antibodies were used and diluted in 10% normal donkey serum: rabbit Ki67 (1:100 Vector laboratories), goat DCX (1:500; Santa Cruz Biotechnology), chicken GFP (1:500; Abcam), rabbit Living Colors^®^ DsRed Polyclonal Antibody (1:1000; Clontech). Neurotrace (Life Technologies) served as a counterstain. All fluorescent secondary antibodies were obtained from Jackson ImmunoResearch and diluted 1:200 in PBS.

As outlined in the Allen Reference atlas(Lein et al., 2007), 20 sagittal sections (S1-S20) of each hemisphere per mouse spaced at 200 μm intervals was analyzed. GFP or Cherry labeled cells in the lateral NDB were counted in sections S13-S15. GFP or Cherry cells in the medial NDB were counted in sections S16-S18. GFP or Cherry cells in the lateral MS were counted in section S20. We did not collect consistent intact samples of section S21 and could not assess the medial MS. For the LC, GFP cells were counted in section S15. LC cells were only detected ipsilateral to the injection. Cherry cells were only detected in the ipsilateral hemisphere. To normalize for viral infection efficiency in each individual mouse, the number of cells in the region of interest (MS, NDB or LC) was divided by the number of cells adjacent to the viral injection site in the dentate gyrus. GFP or Cherry cells were counted in the dentate gyrus, in sections S7 and S8 and averaged. All GFP or Cherry cells in the MS-NDB in sections S13-S20 were assessed for ChAT co-expression and total ChAT cells. DCX and Ki67 were counted unilaterally in 5 sections of dentate gyrus that spanned the septotemporal axis

### Imaging and analysis

Tissue was imaged at 20x on a fluorescent microscope (Olympus IX83). The Allen Brain Atlas was used to define brain regions in sagittal sections. GFP or cherry expressing cells were counted in the medial septum and the diagonal band ipsilateral and contralateral to the injection in sections that transversed the structure.

### Behavioral experiments

All behavioral experiments were performed by a female scientist and took place during the light cycle between 9AM and 3PM. Mice with reduced neurogenesis for 2 months were tested in the following order: spontaneous alternation, open field, fear conditioning. Mice with reduced neurogenesis for 5 months were tested in the following order: spontaneous alternation, open field, elevated plus maze, balance beam. Mice without neurogenesis for 4 months were tested for spontaneous alternation. A group of mice without neurogenesis for 2 months and 12 months were tested in spontaneous alternation and used for pharmacological experiments.

### Spontaneous alternation

To assess spontaneous alternation mice were tested in a closed arm plus maze as described with modifications(Chang and Gold, 2004). The plus maze consisted of four identical arms (25 × 5 × 30 cm) with opaque walls that extended from a center platform (5 × 5 cm) elevated 50 cm from the floor. Testing occurred in a lit (250 lux) room. Mice were placed on the center platform and allowed to explore freely for 12 min. The sequence of arm entries was scored throughout the 12 min. A successful alternation occurred when a mouse made four discrete arm entries on overlapping sets of five consecutive entries. Accordingly the number of successful alternations is the total number of entries minus four. The spontaneous alternation score is calculated by (successful alternations/total possible alternations)x 100; a score of 44% reflects chance performance.

### Open Field

Mice were placed in a Plexiglas open field (Kinder Scientific SmartFrame 22.1” × 22.1” × 15.83”) illuminated by 80-100 lux for thirty minutes. Behavioral measures were automatically recorded by infrared photo beams and analyzed by MotorMonitor software.

### Elevated-plus maze

The elevated plus maze was performed as described(Avgustinovich et al., 2000), and consisted of a central platform (5 × 5 cm) with two opposing open arms (25 × 5 cm) and two opposing arms enclosed by opaque walls (25 × 5 × 30 cm), elevated 50 cm from the floor. Experiments were conducted in the dark with open arms illuminated (100-120 lux). Mice were placed on the central platform facing a closed arm; the number of entries to each arm and duration in each arm was scored for 5 min by an experienced observer.

### Balance Beam

Beam walking was assessed as described(Xie et al., 2010). Mice were given 5 training trials where they traversed a 100 cm long, 1.5 cm diameter circular beam in a lit room (250 lux). 24 hours after training mice traversed the beam once while the number of foot slips and latency to cross the beam were scored.

### Microdialysis

We performed microdialysis to measure baseline and spontaneous alternation induced acetylcholine levels in the dorsal hippocampus as previously described with modifications(Chang and Gold, 2004). NG+ and NG- mice aged without neurogenesis for 1.5 months or 4.5 months were implanted bilaterally with microdialysis guide cannulae (Synaptech, S-3000) in the dorsal hippocampus (bregma coordinates: anteroposterior -2.3mm, mediolateral 1.6mm, dorsoventeral -0.6mm). The cannulae were implanted 1 mm above the dorsal hilus target region as we used a 1 mm membrane to sample from the dorsal hilus. The cannulae were secured to the skull with skull screws and dental cement and mice recovered from surgery for at least 2 weeks.

Microdialysis samples were collected at baseline, during spontaneous alternation and after spontaneous alternation. One group of NG+ and NG- mice were tested after 4 months without neurogenesis with a probe in the left hippocampus. Another group of NG+ and NG- mice were tested after 5 months without neurogenesis with a probe in the right hippocampus and again after 7 months without neurogenesis with a probe in the left hippocampus.

To begin a trial, a microdialysis probe with a 1 mm membrane (Synaptech, S3010 Synaptech Technology Inc., Marquette, MI) was inserted into the dorsal hilus at the same coordinates as the dorsally infused CAV-GFP (bregma coordinates: anteroposterior -2.3mm, mediolateral 1.6mm, dorsoventeral -1.6mm) and mice were placed in an opaque holding cage with fresh bedding. The probes were continuously perfused with 100 nM neostigmine bromide (Sigma) in artificial cerebrospinal fluid (aCSF; 128 mM NaCl, 2.5 mM KCl, 1.3 mM CaCl_2_, 2.1 mM MgCl_2_, 0.9 mM NaH_2_PO_4_, 2.0 mM NaHPO_4_, and 1.0 mM glucose at a pH of 7.4) at 1 μl/min. For the first 60 min mice acclimated to the probe and dialysate was not collected. After this period dialysate samples were collected every 6 min. The first 4 baseline samples were collected while the mouse was in the holding cage. After the baseline sampling time mice were placed in the spontaneous alternation task for 12 min while 2 samples were collected. Finally mice were placed back into the holding cage while 2 post-maze samples were collected.

### HPLC

Dialysate samples were assayed for acetylcholine using HPLC with electrochemical detection (Eicom USA, San Diego, CA). Acetylcholine peaks were quantified by comparison to peak heights of standard solutions and corrected for in vitro recovery of the probe. The system detection limit is reliably 5 femtomole of acetylcholine. Chromatographs obtained every 15 min/sample were analyzed using the software program Envision (provided by Eicom, USA).

### Pharmacology

Scopolamine hydrobromide (Sigma), 0.005 mg/kg dissolved in 0.9% saline was delivered i.p. 40 min prior to the spontaneous alternation task.

Physostigmine hemisulfate (Tocris) 20 ug/kg dissolved in 0.9% saline was delivered i.p. 15 min prior to the spontaneous alternation task.

### Statistics

All statistics were calculated in GraphPad Prism. All data are presented as the means ± SEM and significance was set at *p* < 0.05. ANOVAs that yielded statistically significant main effects were followed with Bonferroni or Tukey’s *post hoc* tests.

